# A general strategy to develop cell permeable and fluorogenic probes for multi-colour nanoscopy

**DOI:** 10.1101/690867

**Authors:** Lu Wang, Mai Tran, Elisa D’Este, Julia Roberti, Birgit Koch, Lin Xue, Kai Johnsson

**Affiliations:** Department of Chemical Biology, Max Planck Institute for Medical Research, Jahnstrasse 29, 69120 Heidelberg, Germany; Optical Microscopy Facility, Max Planck Institute for Medical Research, Jahnstrasse 29, 69120 Heidelberg, Germany; Leica Microsystems CMS GmbH, Am Friedensplatz 3, 68165 Mannheim, Germany; Institute of Chemical Sciences and Engineering, École Polytechnique Fédérale de Lausanne (EPFL), 1015 Lausanne, Switzerland

**Keywords:** Fluorescence, Live-cell imaging, Super-resolution microscopy, Protein labeling, Fluorescent probes

## Abstract

Live-cell fluorescence nanoscopy is a powerful tool to study cellular biology on a molecular scale, yet its use is held back by the paucity of suitable fluorescent probes. Fluorescent probes based on regular fluorophores usually suffer from low cell permeability and unspecific background signal. We report a general strategy to transform regular fluorophores into fluorogenic probes with excellent cell permeability and low unspecific background signal. The strategy is based on the conversion of a carboxyl group found in rhodamines and related fluorophores into an electron-deficient amide. This conversion does not affect the spectroscopic properties of the fluorophore but permits it to exist in a dynamic equilibrium between two different forms: a fluorescent zwitterion and a non-fluorescent, cell permeable spirolactam. Probes based on such fluorophores generally are fluorogenic as the equilibrium shifts towards the fluorescent form when the probe binds to its cellular targets. The resulting increase in fluorescence can be up to 1000-fold. Using this simple design principle we created fluorogenic probes in various colours for different cellular targets for wash-free, multicolour, live-cell nanoscopy. The work establishes a general strategy to develop fluorogenic probes for live-cell bioimaging.

## INTRODUCTION

The combination of fluorescence microscopy and appropriate labeling strategies is a powerful tool to observe cellular structures and biological processes with high spatial and temporal resolution in living cells.^1–3^ However, it has proven to be difficult to design synthetic fluorescent probes suitable for such experiments. A suitable synthetic fluorescent probe for live cell imaging needs to show (i) good cell permeability, (ii) specific staining of its target, (iii) high brightness and photostability, and (iv) suitable excitation/emission wavelengths.^1^ Even though numerous synthetic fluorophores with good photostability and photophysical properties have been developed, most of them endure poor cell permeability.^4, 5^ These properties make them unsuitable for live-cell imaging unless invasive techniques such as permeabilization,^6^ bead loading,^7^ microinjection,^8^ or cell squeezing,^9^ are used to force them into cells. Furthermore, synthetic probes can display high background signal from unspecific binding to intracellular components, requiring time-consuming washing steps.^10^ Fluorogenic probes, i.e. probes that show an increase in fluorescence upon binding to their target, alleviate the problem of background signal. However, the design of such probes is even more challenging.

The majority of fluorescent probes suitable for live-cell microscopy reported so far are based on rhodamine derivatives.^1, 11^ Key to the suitability of rhodamine-based probes for livecell imaging is that they exist in a dynamic equilibrium between a non-fluorescent spirolactone and a fluorescent zwitterion (Fig. 1a). The existence of such a dynamic equilibrium on first sight appears disadvantageous as it reduces the brightness of the fluorophore. However, the hydrophobic, non-fluorescent spirolactone possesses higher cell permeability than its zwitterionic counterpart, while binding of the probe to its target usually shifts the equilibrium towards the fluorescent zwitterion, making such probes fluorogenic. For certain rhodamines such as silicon-rhodamine (SiR)^12^ and JF_585_,^13^ this dynamic equilibrium enables to create highly fluorogenic probes that have become popular tools for live-cell imaging.^12–17^ However, for most rhodamines the formation of the spirolactone is too unfavorable, resulting in low cell-permeability of probes based on these fluorophores. Attempts to shift the equilibrium of such fluorophores more towards the spirolactone are usually based on decreasing the electron density of the xanthene core through introduction of electron-withdrawing groups (EWGs) (blue circles in Figure 1a and Supplementary Fig. S1).^12, 13, 16^ However, conjugation with EWGs may decrease the quantum yield, as is the case with carbopyronines (CPY) and SiR.^16^ Furthermore, reducing the electron density of the xanthene core also increases its susceptibility towards reactions with abundant intracellular nucleophiles.^18–20^

**Figure 1.**
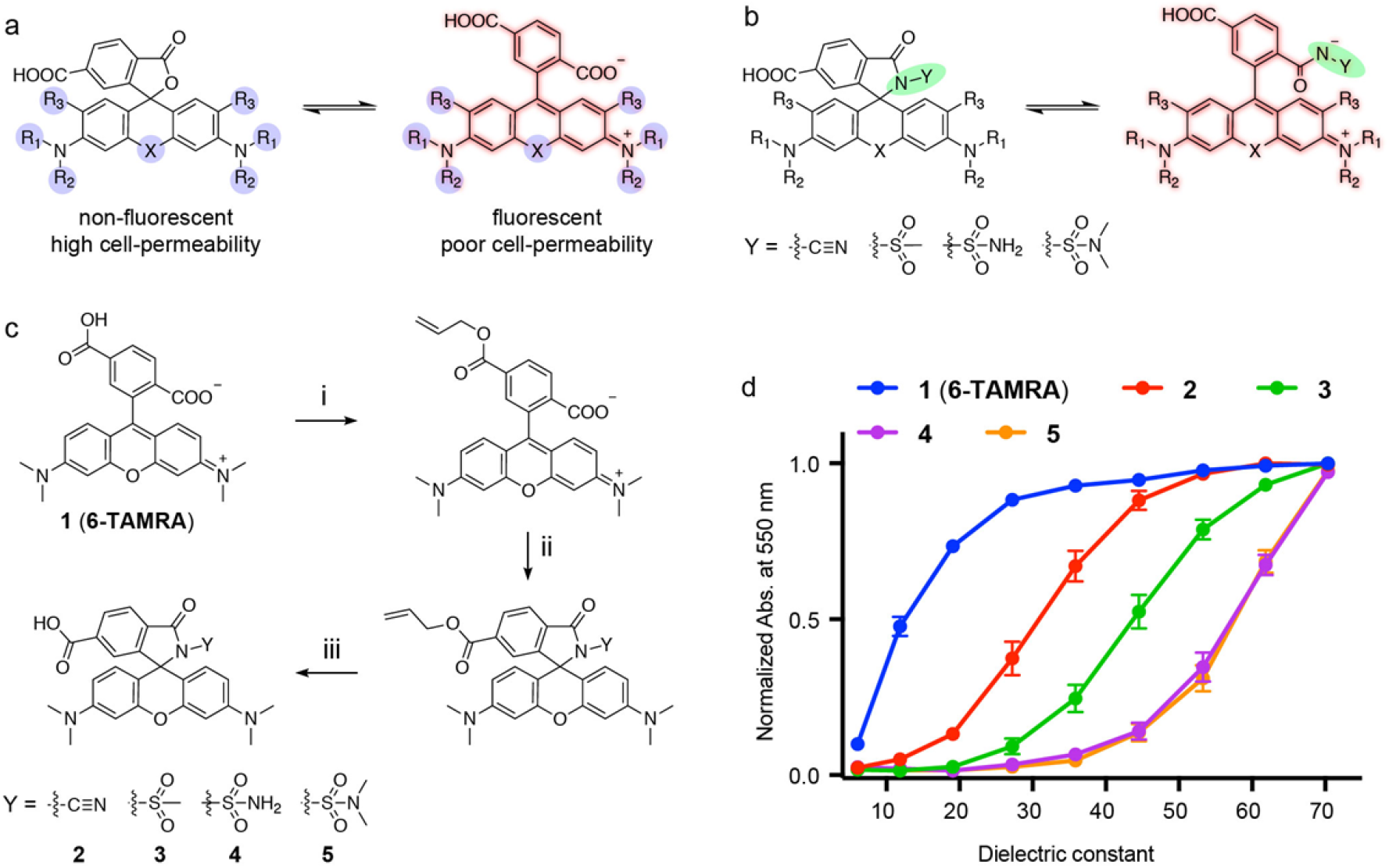
Design strategies for developing cell permeable fluorophores. (**a**) General structure of rhodamines and the equilibrium between the fluorescent zwitterion and the non-fluorescent spirolactone. R_1_-R_3_ (blue circles) denote the positions used for introducing electron-withdrawing groups to favor spirolactone formation. (**b**) General structure of rhodamines and the equilibrium between the fluorescent zwitterion and the non-fluorescent spirolactam. Y (green circle) denotes position for introducing electron-withdrawing groups to disfavor spirolactam formation. (**c**) Synthetic route for preparation of rhodamines (**2-5**). (i) allyl bromide, K_2_CO_3_, Et_3_N, DMF, r.t. 2 h; (ii) POCl_3_, DCM, reflux, 3 h; amines, ACN, DIPEA, r.t. 1 h; (iii) 1,3-Dimethylbarbituric acid/Pd(PPh_3_)_4_, MeOH/DCM, r.t. 1 h. (**d**) Normalized absorbance at 550 nm in zwitterionic form of 5 μM of 6-TAMRA (**1**) and derivatives (**2-5**) in water–dioxane mixtures (v/v: 10/90 - 90/10) as a function of dielectric constant. Error bars show ± s.d. *n* = 3

Here, we introduce a general strategy to increase the cell permeability and fluorogenicity of regular rhodamines without changing their spectroscopic properties. Based on this strategy, we generated rhodamine-based probes (Supplementary Fig. S2) in various colours and with unprecedented fluorogenicity, enabling wash-free, multi-colour live-cell confocal and superresolution microscopy.

## RESULTS AND DISCUSSION

### Designing cell-permeable and fluorogenic rhodamine derivatives

Our efforts to improve the permeability and fluorogenicity of rhodamine derivatives for livecell imaging focused on increasing the propensity of rhodamines to form the spirocyclic, non-fluorescent form without affecting the spectroscopic properties of the parental molecule. We initially considered replacing the carboxylic acid responsible for formation of the spirolactone with an amide (Fig. 1b). However, such rhodamine derivatives have been reported to strongly favor spirolactams at physiological pH,^21, 22^ making them unsuitable for live-cell imaging. We therefore attempted to destabilize the spirolactam by attaching different electron-withdrawing groups to the underlying amides. Specifically, we converted 6-carboxytetramethylrhodamine (6-TAMRA, **1**), a widely used fluorophore with good photophysical properties, into the corresponding acyl cyanamide (**2**), acyl sulfonamide (**3**) and acyl sulfamide (**4** and **5**), (Fig. 1c, Supplementary Table S1 and Scheme S1). The acyl cyanamide of rhodamine B was previously reported to exist as a fluorescent zwitterion at physiological pH.^23^ Acyl sulfonamides and acyl sulfamides are used in medicinal chemistry as anionic pharmacophores.^24, 25^ Amides **2-5** were prepared in three simple steps from commercially available 6-TAMRA (Fig. 1c). To investigate the propensity of these amides to form the corresponding spirolactams, we measured their absorbance spectra in water-dioxane mixtures with different ionic strength. The equilibrium between the zwitterionic and spirocyclic form can then be characterized by the D_50_ value, which is defined as the dielectric constant at which absorbance of the fluorescent zwitterion is decreased by half compared to the highest recorded absorbance value measured in dioxane–water mixtures.^12, 16, 17^ Rhodamine-based fluorophore with D_50_ values around 50 have been shown to be suitable candidates for the generation of fluorogenic probes.^12, 16^ In comparison, the D_50_ value of TAMRA is around 10 showing that in aqueous solution TAMRA exists predominantly in its open form and TAMRA-based probes are usually not fluorogenic. The measured D_50_ values of amides **2-5** ranged from 32 to around 60 (Fig. 1d), which indicates the potential of these fluorophores for the generation of permeable and fluorogenic probes.^16, 17^ Importantly, the spectroscopic properties of amides **2-5** do not differ significantly from 6-TAMRA (Supplementary Table S1). The possibility to tune the equilibrium between the open and closed form of the fluorophore by varying the substituents on the amide is an important feature of this approach.

### Cell-permeable and fluorogenic TAMRA derivatives for no-wash live-cell imaging

Self-labeling protein tags such as SNAP-tag and HaloTag are an efficient approach to attach synthetic fluorophores to proteins in living cells.^26, 27^ The main challenge when using these tags for live-cell imaging is the low cell permeability of most fluorescent probes used for labeling and high background signal resulting from their unspecific binding. SNAP-tag fusion proteins can be specifically labeled with different fluorophores using appropriate O^6^-benzlyguanine derivatives. A 6-TAMRA based probe (**6**, Figure 2a) for SNAP-tag has been previously introduced for live-cell imaging, but its low cell-permeability and lack of fluorogenicity requires the use of high concentrations (5 μM) and repeated washing steps to remove excess and non-specifically bound probe.^28^ To address these problem, we coupled fluorophores **2-5** to O^6^-benzylguanine to obtain probes **7-10** (Fig. 2a). *In vitro* characterization of the spectroscopic properties of these probes in the presence and absence of SNAP-tag showed that in particular probe **10** containing the *N,N*-dimethylsulfamide group (in the following abbreviated as **MaP555-SNAP**) showed a significant increase in absorbance and fluorescence upon binding to SNAP-tag: the absorbance at 555 nm increases 15-fold and the intensity of fluorescence emission at 580 nm increases 21-fold (Fig. 2b and Supplementary Fig. S3). We then examined the performance of probes **6-10** in live-cell imaging, in which U2OS cells stably expressing a nuclear localized SNAP-Halo fusion protein were co-cultured with regular U2OS cells not expressing any protein tags (Figure 2c). The performance of the probes in the *in vitro* tests correlated also with their performance in live-cell imaging: **MaP555-SNAP** even at a very low concentration (250 nM) allowed to perform wash-free labeling with excellent nuclei to cytosol signal ratio (F_nuc._/F_cyt._ = 15), while labeling with control probe **6** under identical conditions only lead to a barely detectable signal (Figure 2c and Supplementary Fig. S4). Furthermore, the kinetics of labeling of nuclear localized SNAP-tag with **MaP555-SNAP** were much faster than those of control probe **6** (Fig. 2d). We then prepared the corresponding 6-TAMRA-based probes **11-15** for HaloTag (Fig. 2e). As for SNAP-tag, the probe with the largest enhancement in absorbance (16-fold) and fluorescence intensity (35-fold) in *in vitro* assays was based on the *N,N*-dimethylsulfamide derivative **15** (abbreviated as **MaP555-Halo** in the following), (Fig. 2f and Supplementary Fig. S5). In no-wash live-cell imaging experiments, **MaP555-Halo** furthermore showed much lower background signal than control probe **11** (Fig. 2g and Supplementary Fig. S6), as most of **MaP555-Halo** exist as the non-fluorescent spirolactam prior to binding to HaloTag. Furthermore, the intracellular kinetics of labeling of nuclear HaloTag with **MaP555-Halo** were about two-fold faster than with control probe **11** (Fig. 2h). The excellent performance of **MaP555-SNAP** and **MaP555-Halo** in live-cell imaging experiments support our assumption that establishing a dynamic equilibrium between the fluorescent zwitterion and the more hydrophobic spirolactam enables the generation of highly cell-permeable and fluorogenic probes for live-cell imaging.

**Figure 2.**
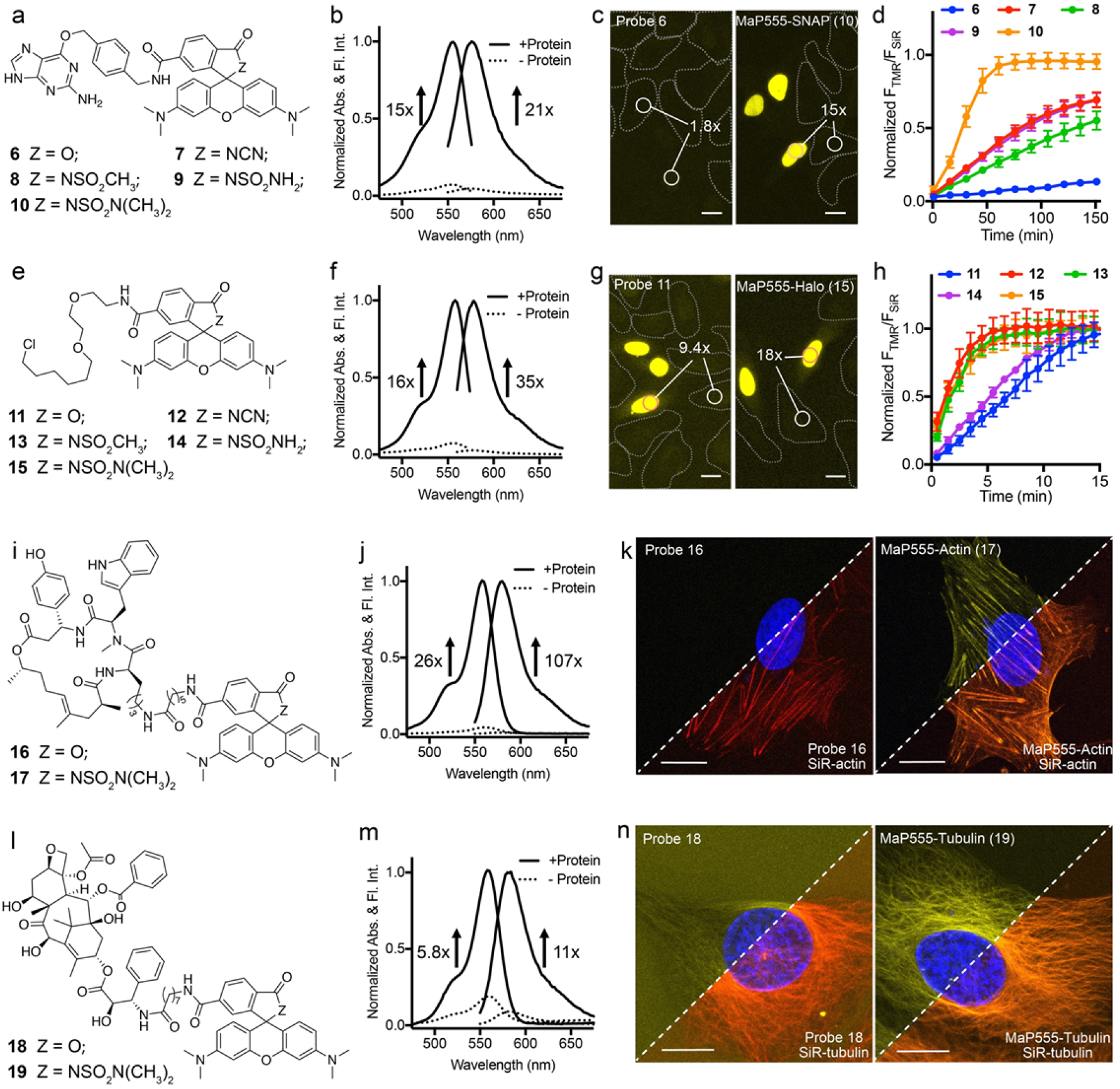
Cell permeable 6-TAMRA derivatives for no-wash live-cell microscopy. **a**, Structures of 6-TAMRA derivatives coupled to SNAP-tag substrate O^6^-benzylguanine (**6-10**). **b**, Absorption and emission spectra of **MaP555-SNAP** (**10**) (2.5 μM) measured in the presence (+protein) and absence (-protein) of SNAP-tag (5 μM) after 1 h incubation. *n* = 3. **c**, No-wash live-cell confocal images of cocultured normal U2OS cells and U2OS FlpIn Halo-SNAP-NLS expressing cells labeled with 250 nM of **6** (left) or **MaP555-SNAP** (**10**) (right) for 1.5 h. **d**, Normalized ratio of TAMRA fluorescence (**6-10**) to SiR fluorescence at different time points in U2OS FlpIn Halo-SNAP-NLS expressing cells. The cells were pre-labeled with 500 nM of SiR-Halo for 2 h and then treated with 250 nM of **6-10**. *n* > 150 cells. e, Structures of 6-TAMRA derivatives coupled to HaloTag substrate (**11-15**). **f**, Absorption and emission spectra of **MaP555-Halo** (**15**) (2.5 μM) measured in the presence (+protein) and absence (protein) of HaloTag (5 μM) after 1 h incubation. *n* = 3. **g**, No-wash, live-cell confocal images of cocultured normal U2OS cells and U2OS FlpIn Halo-SNAP-NLS expressing cells labeled with 100 nM of **11** (left) or **MaP555-Halo** (**15**) (right) for 30 min. h, Normalized ratio of TAMRA fluorescence (**11-15**) to SiR fluorescence at different time points in U2OS FlpIn Halo-SNAP-NLS expressing cells. The cells were pre-labeled with 500 nM of SiR-BG for 2 h and then treated with 100 nM of 11-15. *n* > 150 cells. i, Structures of actin probe 16 and **MaP555-actin** (17). **j**, Absorption and emission spectra of **MaP555-actin** (17) (2.5 μM) measured in the presence (+protein) and absence (-protein) of actin (0.4 mg/mL) after 2 h incubation. *n* = 3. **k**, Images of no-wash live U2OS cells labeled with 500 nM of **16** (left) or **MaP555-actin** (17) (right) and 500 nM of **SiR-actin** for 1 h. Single focus plane. l, Structures of tubulin probe **18** and **MaP555-tubulin** (19). m, Absorption and emission spectra of **MaP555-tubulin** (**19**) (2.5 μM) measured in the presence (+protein) and absence (-protein) of tubulin (2 mg/mL) after 2 h incubation. *n* = 3. n, No-wash images of live U2OS cells labeled with 1 μM of **18** (left) or **MaP555-tubulin** (**19**) (right), 1 μM **SiR-tubulin** and 0.5 μM of verapamil for 1.5 h. Scale bar (**c, g, k, n**), 20 μm. Normal U2OS cells in **c** and **g** were marked with white dashed lines. Numbers in c and g indicate the average ratios between nuclear signal (U2OS FlpIn Halo-SNAP-NLS expressing cells) and cytosol signal (normal U2OS cells) (F_nuc._/F_cyt._). *n* (**c, g**) > 200 cells. Error bars show ± s.e.m.

Fluorescent probes for live-cell imaging of cytoskeletal structures have become important tools in the life sciences.^1^ In particular, the far-red, SiR-based probes for F-actin (**SiR-actin**) and microtubules (**SiR-tubulin**) have become popular as they are fluorogenic and enable no-wash imaging with little background signal.^14^ **SiR-actin** and **SiR-tubulin** are based on SiR linked to the F-actin-binder jasplakinolide and microtubule-binder docetaxel, respectively. Binding to their targets shifts for both probes the equilibrium between the cell-permeable spirolactone and the fluorescent zwitterion towards the fluorescent form. To generate fluorescent stains for F-actin and microtubule in different colors, we coupled jasplakinolide and docetaxel to 6-TAMRA (probes **16** and **18**, Figure 2i, l) and its *N,N*-dimethylsulfamide derivative (probes **17** and **19**, named **MaP555-actin** and **MaP555-tubulin** in the following, Figure 2i, l). As expected,^29^ the 6-TAMRA derivatives of jasplakinolide 16 and docetaxel 18 did not allow to perform live-cell imaging of F-actin and microtubules in U2OS cells (Fig. 2k, n), presumably because these probes predominantly exist as zwitterions with low permeability. In contrast, **MaP555-actin** and **MaP555-tubulin** enabled no-wash, high-contrast staining of F-actin and microtubules in U2OS cells (Fig. 2k, n). The specificity of **MaP555-actin** and **MaP555-tubulin** was further confirmed by co-staining with **SiR-actin** and SiR-tubulin (Fig. 2k, n). The performance of both probes can be rationalized by their fluorogenicity: In *in vitro* assays, **MaP555-actin** showed a large increase in absorbance (26-fold) and fluorescence intensity (107-fold) upon binding to F-actin (Fig. 2j and Supplementary Fig. S7). Similarly, binding of **MaP555-tubulin** to microtubules caused a high increase in absorbance (5.8-fold) and fluorescence (11-fold) (Fig. 2m and Supplementary Fig. S7).

### Extension of the strategy to other rhodamine derivatives

Encouraged by the outstanding performance of these new TAMRA-based probes, we extended our design strategy to other commonly used rhodamine derivatives with wavelengths ranging from cyan to near-infrared. Rhodamine 110 (R110) is a classic fluorophore emitting cyan fluorescence with good photostability and photophysical properties.^22^ However, its low D_50_ value of 15 and the presence of two polar NH_2_ groups results in low cell permeability of R110-based probes. Live-cell imaging of R110-based probes thus often requires high concentrations, long incubation times, and tedious washing steps.^13, 30, 31^ The introduction of *N,N*-dimethylsulfamide into R110 (**20**) to yield fluorophore **MaP510** (**21**) shifted its D_50_ from 15 to 70 (Supplementary Fig. S8). The corresponding probe for HaloTag, **MaP510-Halo** (**23**, Fig. 3a) showed a large increase response in absorbance and fluorescence (14-fold and 11-fold, respectively) upon binding to HaloTag *in vitro* (Fig. 3b, Supplementary Fig. S9, and Table S2). In contrast, the regular R110-based HaloTag probe **22** showed no significant fluorogenicity (Supplementary Fig. S9). In addition, **MaP510-Halo** even at a low concentration (250 nM) resulted in fast staining (< 30 min) of nuclear localized HaloTag in U2OS cells with high signal to background ratio (18-fold), whereas control compound **22** showed a much lower signal to background ratio (2.5-fold) under these conditions (Fig. 3c and Supplementary Fig. S10).

**Figure 3.**
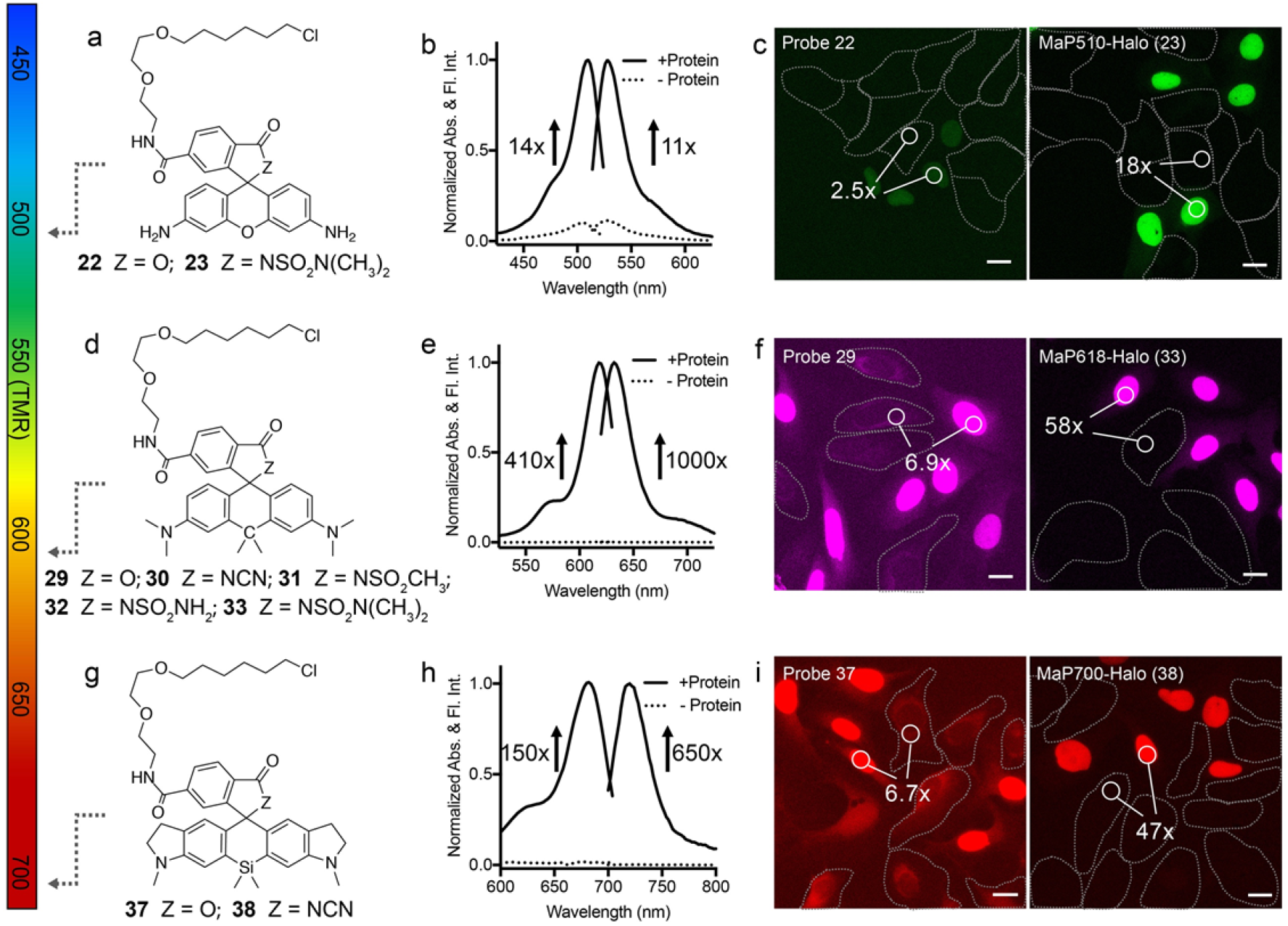
Cell permeable probes with wavelengths ranging from cyan to NIR for no-wash, livecell microscopy. **a**, Structures of R110 derivatives coupled to HaloTag substrate (**22** and **MaP510-Halo** (**23**)). **b**, Absorption and emission spectra of **MaP510-Halo** (**23**) (2.5 μM) measured in the presence (+protein) and absence (-protein) of HaloTag (5 μM) after 1 h incubation. *n* = 3. **c**, No-wash live-cell confocal images of co-cultured normal U2OS cells and U2OS FlpIn Halo-SNAP-NLS expressing cells labeled with 250 nM of **22** (left) or **MaP510-Halo** (**23**) (right) for 30 min. **d**, Structures of CPY derivatives coupled to HaloTag substrate (**29-33**). **e**, Absorption and emission spectra of **MaP618-Halo** (**33**) (2.5 μM) measured in the presence (+protein) and absence (-protein) of HaloTag (5 μM) after 1 h incubation. *n* = 3. **f**, No-wash, live-cell confocal images of co-cultured normal U2OS cells and U2OS FlpIn Halo-SNAP-NLS expressing cells labeled with 500 nM of 29 or **MaP618-Halo** (**33**) for 30 min. **g**, Structures of SiR700 derivatives with HaloTag substrate (37 and **MaP700-Halo** (**38**)). **h**, Absorption and emission spectra of **MaP700-Halo** (38) (2.5 μM) measured in the presence (+protein) and absence (-protein) of HaloTag (5 μM) after 1 h incubation. *n* = 3. **i**, No-wash live-cell confocal images of co-cultured normal U2OS cells and U2OS FlpIn Halo-SNAP-NLS expressing cells labeled with 250 nM of 37 or **MaP700-Halo** (**38**) for 30 min. Scale bar (*c, f, i*), 20 μm. Normal U2OS cells were marked with white dashed lines. Numbers in **c, f**, and **i** indicate the average ratios between nuclear signal (U2OS FlpIn Halo-SNAP-NLS expressing cells) and cytosol signal (normal U2OS cell) (Fnuc./Fcyt.). *n* (**c, f, i**) > 200 cells.

We next applied this strategy to carbopyronine, an orange fluorophore whose brightness and photostability make it an attractive choice for confocal and superresolution microscopy.^16^ Introduction of the cyanamide, sulfonamide and the two sulfamides into carbopyronine (Supplementary Fig. S11), allowed us to generate HaloTag probes **30-33** (Fig. 3d). All four probes showed large increases in fluorescence intensities upon binding to HaloTag, with values ranging from 100-to a 1000-fold (Fig. 3e, Supplementary Fig. S12 and Table S3). In contrast, the regular carbopyronine HaloTag probe **29** only showed a 3.8-fold increase in fluorescence upon HaloTag binding (Supplementary Fig. S12). The observed 1000-fold increase in fluorescence intensity of probe **33** (in the following named **MaP618-Halo**) and its brightness sets this probe apart from other fluorogenic HaloTag substrates reported previously.^12, 13, 32^ The outstanding fluorogenicity of **MaP618-Halo** and the other carbopyronine-based probes 30-33 make them powerful probe for live-cell imaging: no-wash live-cell imaging of U2OS cells expressing nuclear localized HaloTag showed bright nuclei with extremely low unspecific extranuclear fluorescence (F_nuc._/F_cyt._ = 58) (Fig. 3f and Supplementary Fig. S13). In contrast, labeling with the regular carboypyronine probe **29** resulted in significantly higher unspecific extranuclear fluorescence (F_nuc._/F_cyt._ = 6.8). Furthermore, labeling with **MaP618-Halo** was very rapid and reached saturation within 5 min (Supplementary Fig. S14).

In addition, we coupled the acyl sulfonamide of carbopyronine **26** to jasplakinolide, yielding the actin probe **MaP618-actin** (**34**). **MaP618-actin** possesses orange fluorescence (***λ***_ex_/***λ***_em_: 618/635 nm) (Supplementary Fig. S15) and its absorbance and fluorescence intensity increase 122-fold and 449-fold upon incubation with F-actin (Supplementary Fig. S15). **MaP618-actin** is thus about four times more fluorogenic than the previously described **SiR-actin**. Micrographs of live U2OS cells incubated with 500 nM of **MaP618-actin** for 1 h without any washing steps clearly reveal F-actin structures, which were also verified by colocalization with **SiR-actin** (Supplementary Fig. S16). Overall, these data clearly demonstrate the potential of **MaP618-actin** for live-cell imaging of F-actin.

Fluorophores with absorption and emission wavelength in the NIR windows are attractive choices for live-cell and *in vivo* imaging because of the reduced autofluorescence, deep tissue penetration and decreased phototoxicity at this wavelength.^33, 34^ We have previously described the NIR probe silicon-rhodamine 700 (SiR700, **35**) (Supplementary Fig. S17). SiR700 possesses fluorogenic properties and SiR700-based probes have been successfully used for live-cell imaging of microtubule, F-actin and lysosomes in no-wash live-cell imaging.^17^ However, a SiR700-based probe for HaloTag (**37**, Fig. 3g) showed only modest fluorogenicity and relatively high background signal in live-cell imaging (Fig. 3i and Supplementary Fig. S18). To solve this problem, we incorporated the acyl cyanamide into SiR700 (**36**, Supplementary Fig. S18). **36**, which exists predominantly as the spirolactam in aqueous solution (Supplementary Fig. S17), was then used to prepare a probe for HaloTag (**MaP700-Halo**, 38) (Fig. 3g and Supplementary Table S4). Incubation of **MaP700-Halo** with HaloTag resulted in a dramatic increase in absorbance (150-fold) and fluorescence intensity (650-fold) (Fig. 3h). Most importantly, U2OS cells incubated with 250 nM of **MaP700-Halo** and imaged without any washing steps displayed bright nuclear fluorescence and negligible background (F_nuc._/F_cyt._ =47, Fig. 3i). Furthermore, labeling of nuclear localized HaloTag in U2OS cells with **MaP700-Halo** (250 nM) was completed within 10 min (Supplementary Fig. S19). These features make **MaP700-Halo** an appealing probe for live-cell and *in vivo* imaging.

The experiments described above demonstrate how the introduction of acyl amides into rhodamine derivatives can be used to dramatically increase both their fluorogenicity and cell-permeability. The availability of different acyl amides that vary in their propensities for spirolactam formation facilitates for a given target the design of fluorescent probe that has the right balance between cell permeability, fluorogenicity and brightness.

### Applications in no-wash, multicolour confocal and STED microscopy

Mechanistic studies of most biological processes require the simultaneous imaging of multiple biomolecules and biochemical activities. Synthetic fluorescent probes for multicolour, live-cell imaging need to be spectrally distinguishable, cell-permeable and ideally should be suitable for no-wash imaging. However, due to the paucity of suitable fluorescent probes that fulfil these requirements, very few no-wash, multicolour, live-cell imaging experiments have been reported so far.^17^ The cell-permeability and fluorogenicity of the here introduced **MaP510, MaP555, MaP618** and **MaP700** probes make them attractive candidates for no-wash, multicolour microscopy. In proof-of-principle experiments, U2OS cells stably expressing mitochondrial localized Cox8-Halo-SNAP were incubated with Hoechst (0.2 μg/mL), **MaP555-tubulin** (1 μM), **MaP618-actin** (500 nM), and **MaP700-Halo** (250 nM) for 1.5 h and imaged directly without any washing steps (Fig. 4a). In addition, time-lapse movies enabled us to follow dynamic changes of the labelled structures (Supplementary Video 1). To underscore the potential of our **MaP** probes for multicolour imaging, various other combinations of fluorescent probes for no-wash, live-cell confocal imaging were successfully tested (Fig. 4b, Supplementary Fig. S20 and Supplementary Videos 2 and 3).

**Figure 4.**
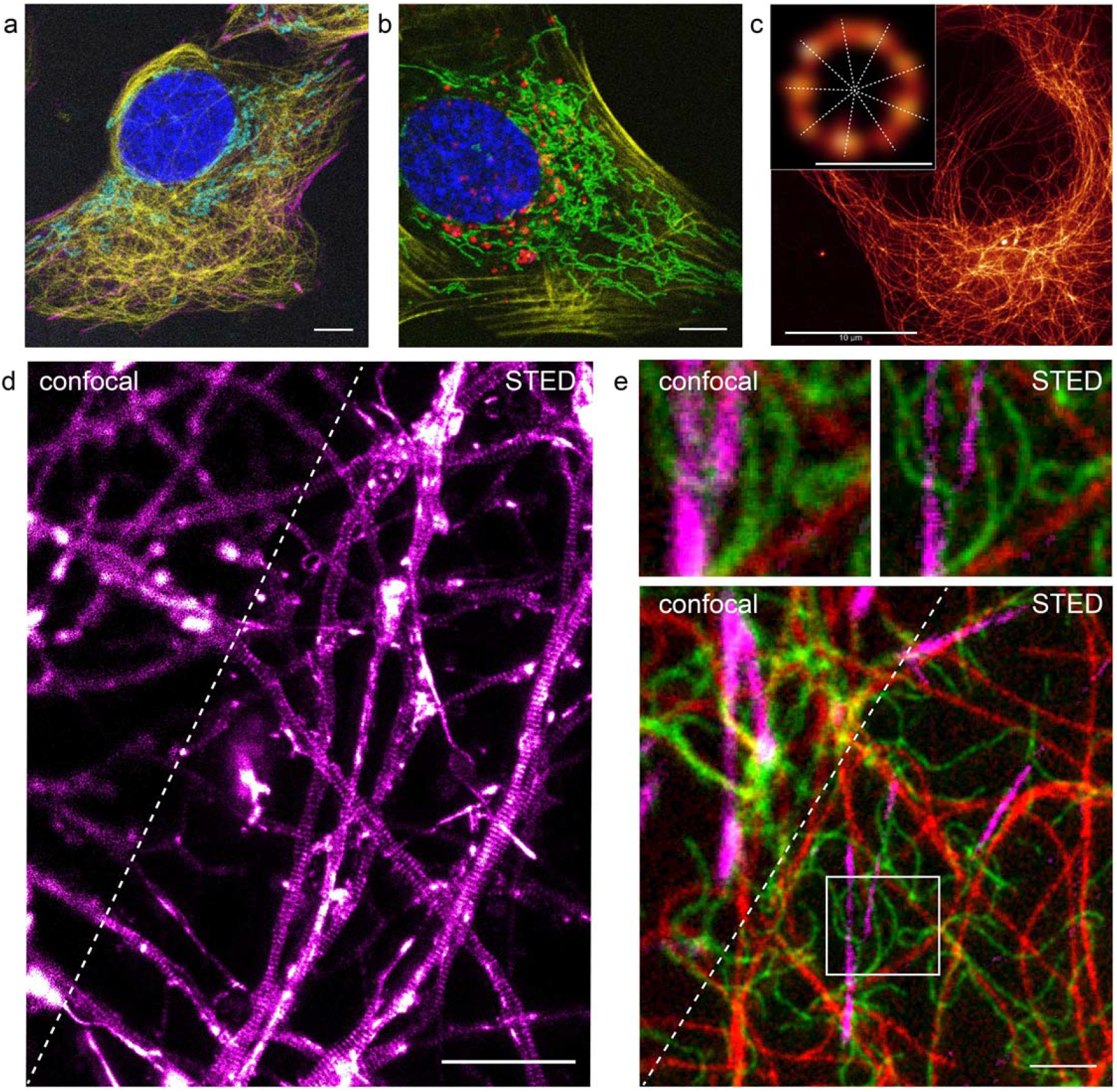
No-wash live-cell confocal and STED microscopy. **a**, No-wash four-colour confocal image of live U2OS FlpIn Cox8-Halo-SNAP expressing cells stained with 0.2 μg/mL of Hoechst (blue), 1 μM of **MaP555-tubulin** (yellow), 500 nM of **MaP618-actin** (magenta), 250 nM of **MaP700-Halo** (cyan), and 5 μM of verapamil for 1.5 h. **b**, No-wash four-colour confocal image of live U2OS FlpIn Cox8-Halo-SNAP expressing cells stained with 0.2 μg/mL of **Hoechst** (blue), 250 nM of **MaP510-Halo** (green), 1 μM of **MaP555-actin** (yellow), and 1 μM of **SiR-Lyso** (red) for 1 h. Scale bar, 10 μm (**a, b**). **c**, No-wash STED images showing the microtubules in live U2OS cells labeled with 1 μM of **MaP555-tubulin** for 2 h. Scale bar, 10 μm. Insert (top left) shows deconvoluted image of selected centriole; the nine-intensity maxima representing the nine microtubule triples are depicted by dashed lines. Scale bar, 200 nm. **d**, No-wash confocal and STED images (raw data) showing actin periodic structures in the axons of rat primary hippocampal neurons stained with 500 nM of **MaP618-actin** for 1.5 h. Scale bar, 5 μm. **e**, No-wash, three-colour confocal and STED images of live U2OS Vimentin-Halo expressing cells stained with 500 nM of **MaP510-Halo** (green, STED at 595 nm), 1 μM of **MaP555-tubulin** (red, STED at 775 nm), 1 μM of **MaP618-actin** (magenta, STED at 775 nm), and 10 μM of verapamil for 2 h. Images (top) shows zoom-in confocal and STED images of the region shown in panel as a white rectangle. Image data were smoothed with a 1-pixel low pass Gaussian filter. Scale bar, 2 μm.

Stimulated Emission Depletion (STED) microscopy is a powerful tool to image biological structures in living cells on the nanoscale. As for conventional microscopy, the impact of livecell STED nanoscopy is limited by the number of available fluorescent probes. The spectroscopic properties of the fluorophores on which our **MaP** probes are based are all compatible with STED nanoscopy and we therefore investigated their performance in such experiments. We first utilized **MaP555-tubulin** to image microtubules in live U2OS cells, using a 660 nm STED beam. In these experiments, U2OS cells were incubated with 1 μM of **MaP555-tubulin** for 1 h and subsequently imaged without any washing steps. The images showed both peripheral microtubules and the microtubules of the centrosomes (Fig. 4c and Supplementary Fig. S21). The apparent diameter of peripheral microtubules determined in these experiments was 39.5 ± 10 nm, a value which is similar to the one obtained with SiR-tubulin^14^. These images also provided a detailed view of the structure of the centriole, the structure around which the centrosome is assembled. Centrioles are cylindrical structures composed of nine triplets of microtubules, with the triplets forming the outer ring of the cylinder.^35^ The observed intensity maxima along the ring of the cylinder clearly revealed the nine-fold symmetry of the centriole. To our knowledge, it is the first time that the individual microtubule triplets are resolved in no-wash, live-cell imaging experiments. Similar to **MaP555-tubulin**, **MaP555-actin** also allowed to image the actin cytoskeleton in U2OS cells with nanoscale resolution (Supplementary Fig. S21).

We previously used live-cell STED nanoscopy and **SiR-actin** to image the periodic arrangement of the actin subcortical cytoskeleton along the neurites of hippocampal neurons.^36^ To increase the signal-to-noise ratio in these experiments, excess probe needs to be removed through a washing step. Using **MaP618-actin**, we were able to directly image this subcortical actin structure under no-wash conditions (Fig. 4d and Supplementary Fig. S22). We attribute the increased signal-to-noise ratio of **MaP618-actin** relative to that of **SiR-actin** to its increased fluorogenicity.

Having a set of spectrally orthogonal and highly fluorogenic probes at hand, we next tested their performance in no-wash, multicolour, live-cell STED nanoscopy. U2OS stably expressing vimentin-HaloTag were incubated with **MaP510-Halo** (green), **MaP555-tubulin** (red), and **MaP618-actin** (magenta) for 2 h and imaged directly. In these experiments, **MaP555-tubulin** and **MaP618-actin** were simultaneously imaged using a 775 nm depletion laser. Subsequently, **MaP510-Halo** was imaged using a 595 nm depletion laser. In this way, three-color, no-wash STED images at sub-diffraction resolution and without any cross-talk between the different channels were obtained (Fig. 4e and Supplementary Fig. S23).

Overall, these experiments clearly highlight the potential of the here introduced probes for live-cell nanoscopy.

## CONCLUSION

We have introduced a general strategy to generate fluorogenic probes with excellent cell permeability and low unspecific background signal. Using this strategy, we created probes with wavelengths ranging from cyan to near-infrared for SNAP-tag, HaloTag, F-actin and microtubule. The good spectroscopic properties, high cell permeability and outstanding fluorogenicity of these probes will make them powerful tools for live-cell nanoscopy. Furthermore, the generality and versatility of the established design principle can be employed to improve the performance of many other already existing fluorescent probes and opens up the opportunity to create numerous new ones.

## METHODS

Detailed procedures for the synthesis of all compounds, their characterization and imaging experiments are given in the Supplementary Information.

### Live-cell labelling and single colour confocal imaging

The Flp-In™ System (ThermoFisher Scientific) was used to generate U2OS FlpIn Halo-SNAP-NLS expressing cells. Briefly, pcDNA5-FRT-Halo-SNAP-NLS and pOG44 were co-transfected into the host cell line U2OS FLpIn.^37^ Homologous recombination between the FRT sites in pcDNA5-FRT-Halo-SNAP-NLS and on the host cell chromosome, catalyzed by the Flp recombinase expressed from pOG44, produced the U2OS FlpIn Halo-SNAP-NLS stably expressing cells. Live U2OS FlpIn Halo-SNAP-NLS expressing cells were labelled and imaged in imaging medium containing phenol red free Dulbecco’s Modified Eagle Medium (DMEM) media that contained 10% FBS, 1 mM pyruvate, and 2 mM L-Glutamine (Thermo Fisher Scientific). Live U2OS cells were incubated with 0.1 to 1 μM of probes for 0.5 to 2 h at 37 °C in a 5% CO_2_ atmosphere. The cells were then directly imaged using confocal microscopy without washing steps. A Leica SP8 confocal scanning microscope equipped with a white light laser, HC PL APO CS2 20.0x objective lens and spectral HyD detector was used. Microscopy conditions: R110 derivatives, ***λ***_ex_: 505 nm, detection range: 515 − 600 nm; TAMRA derivatives, ***λ***_ex_: 555 nm, detection range: 565 − 650 nm; CPY derivatives, ***λ***_ex_: 605 nm, detection range: 615 − 700 nm; SiR700 derivatives, ***λ***_ex_: 670 nm, detection range: 680 – 780 nm. Images are presented as maximum intensity projections (z-stack, 20 slices, 25 μm) unless otherwise noted.

### No wash live-cell multi-colour confocal microscopy

U2OS FlpIn Halo-SNAP-NLS and U2OS FlpIn Cox8-Halo-SNAP expressing cells were grown in 96-well plate (Thermos Fisher) with optical bottom at 37 °C in a humidified 5% CO_2_ environment. Probes (0.1 - 1 μM) were added and the cells were incubated for 1 - 2 h in the above imaging medium. 5 μM of verapamil was added when staining with **MaP555-tubulin**. The cells were directly imaged on a single focal plane. A Leica SP8 microscope equipped with a white light laser and HC PL APO CS2 40.0x/1.10 water objective was used. Images were obtained with sequence program in Leica LAS X software. Microscopy conditions: Hoechst 33343: ***λ***_ex_: 405 nm, detection range: 415 – 470 nm; R110 derivatives, detection range: 505 nm, filter: 515 – 560 nm; TAMRA derivatives, ***λ***_ex_: 555 nm, detection range: 565 – 625 nm; CPY derivatives, ***λ***_ex_: 595 nm, detection range: 605 – 645 nm; SiR derivatives, ***λ***_ex_: 640 nm, detection range: 655 – 715 nm; SiR700 derivatives, ***λ***_ex_: 670 nm, detection range: 695 – 735 nm.

### No wash live-cell STED microscopy

U2OS Vimentin-Halo cells were seeded on glass coverslips and were incubated in imaging medium that contained 0.5 μM of **MaP510-Halo**, 1 μM of **MaP555-tubulin**, 1 μM of **MaP618-actin** and 10 μM of Verapamil for 2 h at 37 °C. Then the live cells were imaged on a Abberior STED 775/595/RESOLFT QUAD scanning microscope (Abberior Instruments GmbH) equipped with STED lines at 595 and 775 nm, excitation lines at 355 nm, 405 nm, 485 nm, 561 nm, and 640 nm, spectral detection, and a UPlanSApo 100x/1.4 oil immersion objective lens. The imaging conditions are listed below: **MaP510-Halo** (***λ***_ex_: 485 nm, detection range: 505-550 nm, STED Laser: 595 nm); **MaP555-tubulin** (***λ***_ex_: 561 nm, detection range: 575-605 nm, STED Laser: 775 nm); **MaP618-actin** (***λ***_ex_: 640 nm, detection range: 655-700 nm, STED Laser: 775 nm).

Primary hippocampal neurons were prepared from postnatal P0-P2 Wistar rats as previously described^14^ and cultured on glass coverslips for 27 days. The procedure was conducted in accordance with Animal Welfare Law of the Federal Republic of Germany (Tierschutzgesetz der Bundesrepublik Deutschland, TierSchG) and the regulation about animals used in experiments (August 1, 2013, Tierschutzversuchsverordnung). For the procedure of euthanizing rodents for subsequent preparation of any tissue, all regulations given in §4 TierSchG are followed. Because euthanization of animals is not an experiment on animals according to §7 Abs. 2 Satz 3 TierSchG, no specific authorization or notification is required. Neuron cells were incubated with 0.5 μM of **MaP618-actin** in ACSF (artificial cerebrospinal fluid) for 1.5 h at 37 °C and directly imaged with the above STED conditions on a single focal plane.

U2OS cells seeded on 8-well chambers (IBIDI) were incubated in imaging medium that contained 1 μM of **MaP555-tubulin** or 1 μM of **MaP555-actin** for 2 h at 37 °C. Then, the live cells were imaged on a Leica TCS SP8 STED 3X microscope (Leica Microsystems GmbH) equipped with STED lines at 592, 660 and 775 nm, excitation with a supercontinuum White Light Laser (WLL) source, spectral detection, and a STED WHITE 86x/1.2 W water immersion objective lens. The imaging conditions were optimized for STED performance of the TAMRA derivatives: ***λ***_ex_: 555 nm corresponding to the maximum of the excitation spectrum, detection range: 565-650 nm, STED Laser: 660 nm. Adaptive deconvolution for centriole was applied using the Leica LIGHTNING detection package (Leica Microsystems GmbH).

## Acknowledgements

The authors acknowledge funding from the Max Planck Society. We thank Prof. Dr. Stefan Jakobs (Max-Planck-Institute for Biophysical Chemistry, Göttingen) for kindly providing the U2OS Vimentin-HaloTag cells. Jasmine Hubrich and Clara-Marie Gürth supported with cell culture and preparation of neurons. We are grateful to Stefan Pitsch (Spirochrome A.G.) for the gift of SiR700 and to Luc Reymond (EPFL) for the gift of carbopyronine. L.W. and M.T. were supported by fellowships of the Alexander von Humboldt Foundation.

## Author contributions

All authors planned the experiments and contributed to paper writing. L.W. and K.J. designed the strategy and fluorophore structures. L.W., M.T. and L.X performed the chemical syntheses. L.W. and M.T. characterized the dyes. L.W. and M.T. performed confocal microscopy with subsequent data analysis. E.D’E. and J.R. performed STED microscopy with subsequent data analysis. B.K. developed cell lines.

## Additional information

Supplementary information and chemical compound information are available in the online version of the paper. Reprints and permission information is available online at http://www.nature.com/reprints. Correspondence and requests for materials should be addressed to K.J.

